# Neuronal oscillatory activities in separate frequencies encode hierarchically distinct visual features

**DOI:** 10.1101/2020.01.13.902775

**Authors:** Hiroto Date, Keisuke Kawasaki, Isao Hasegawa, Takayuki Okatani

**Affiliations:** Graduate School of Information Sciences, Tohoku University; Graduate School of Medical and Dental Sciences, Niigata University; Center for Advanced Intelligence Project, Riken

## Abstract

Although most previous studies in cognitive neuroscience have focused on the change of the neuronal firing rate under various conditions, there has been increasing evidence that indicates the importance of neuronal oscillatory activities in cognition. In the visual cortex, specific time-frequency bands are thought to have selectivity to visual stimuli. Furthermore, several recent studies have shown that several time-frequency bands are related to frequency-specific feedforward or feedback processing in inter-areal communication. However, few studies have investigated detailed visual selectivity of each time-frequency band, especially in the primate inferior temporal cortex (ITC). In this work, we analyze frequency-specific electrocorticography (ECoG) activities in the primate ITC by training encoding models that predict frequency-specific amplitude from hierarchical visual features extracted from a deep convolutional neural network (CNNs). We find that ECoG activities in two specific time-frequency bands, theta (around 5 Hz) and gamma (around 20-25 Hz) bands, are better predicted from CNN features than the other bands. Furthermore, theta- and gamma-band activities are better predicted from higher and lower layers in CNNs, respectively. Our visualization analysis using CNN-based encoding models qualitatively show that theta- and gamma-band encoding models have selectivity to higher- and lower-level visual features, respectively. Our results suggest that neuronal oscillatory activities in theta and gamma bands carry distinct information in the hierarchy of visual features, and that distinct levels of visual information are multiplexed in frequency-specific brain signals.

## 1 Introduction

One important goal in cognitive neuroscience is to understand the relationship between brain activities and sensory information. From single-neuron spikes to mesoscopic brain signals, brain activities can be measured in a wide range of scales of the brain. In the literature, most previous studies have investigated the change of the neuronal firing rate to various synthetic or natural stimuli. However, there has been increasing evidence that indicates the importance of neuronal oscillatory activities in cognition [1, 2].

Neuronal oscillatory activities can be measured by local field potentials (LFPs), electrocorticography (ECoG), electroencephalography (EEG), and magnetoencephalography (MEG). It is thought that raw brain signals contain aggregated activities in several time-frequency bands, such as delta, theta, alpha, beta, and gamma bands [3]. In the visual cortex, several previous studies suggest that brain signals in specific time-frequency bands have stronger selectivity to visual stimuli [4, 5, 6, 7], and that low- and high-frequency bands are related to distinct visual information [8, 9, 10]. Furthermore, different frequency bands are thought to play complementary roles in inter-areal feedforward and feedback processing [11, 12, 13, 14]. However, few studies have investigated the detailed visual selectivity of each frequency band, especially in the primate inferior temporal cortex (ITC), which is the highest-level area in the ventral visual pathway.

In this work, to investigate how visual selectivity differs across frequency bands, we analyze frequency-specific activities in ECoG signals recorded from the macaque ITC using rich hierarchical visual representations extracted from a deep convolutional neural network (CNN). CNNs have achieved state-of-the-art performance on various computer vision tasks [15, 16, 17, 18]. Furthermore, CNNs enable us to extract optimized hierarchical visual features. Previous studies indicate that units in lower, mid, and higher CNN layers represent lower-, mid-, and higher-level visual features, respectively [19]. Several recent studies in neuroscience used visual features in CNNs for analyzing the similarity of the layer hierarchy of CNNs and the anatomical hierarchy of the ventral visual pathway [20, 21]. In our experiments, we trained and evaluated encoding models that predict frequency-specific ECoG activities from visual features extracted at a specific layer of a pretrained CNN (Figure 1). We found that two specific frequency bands, theta (around 5 Hz) and gamma (around 20-25 Hz) bands, were better predicted from CNN features than the other bands. Furthermore, these two bands were better predicted from higher and lower CNN layers, respectively. Our visualization analysis using CNN-based encoding models qualitatively showed that theta- and gamma-band encoding models had selectivity to higher- and lower-level visual features, respectively.

**Figure 1:**
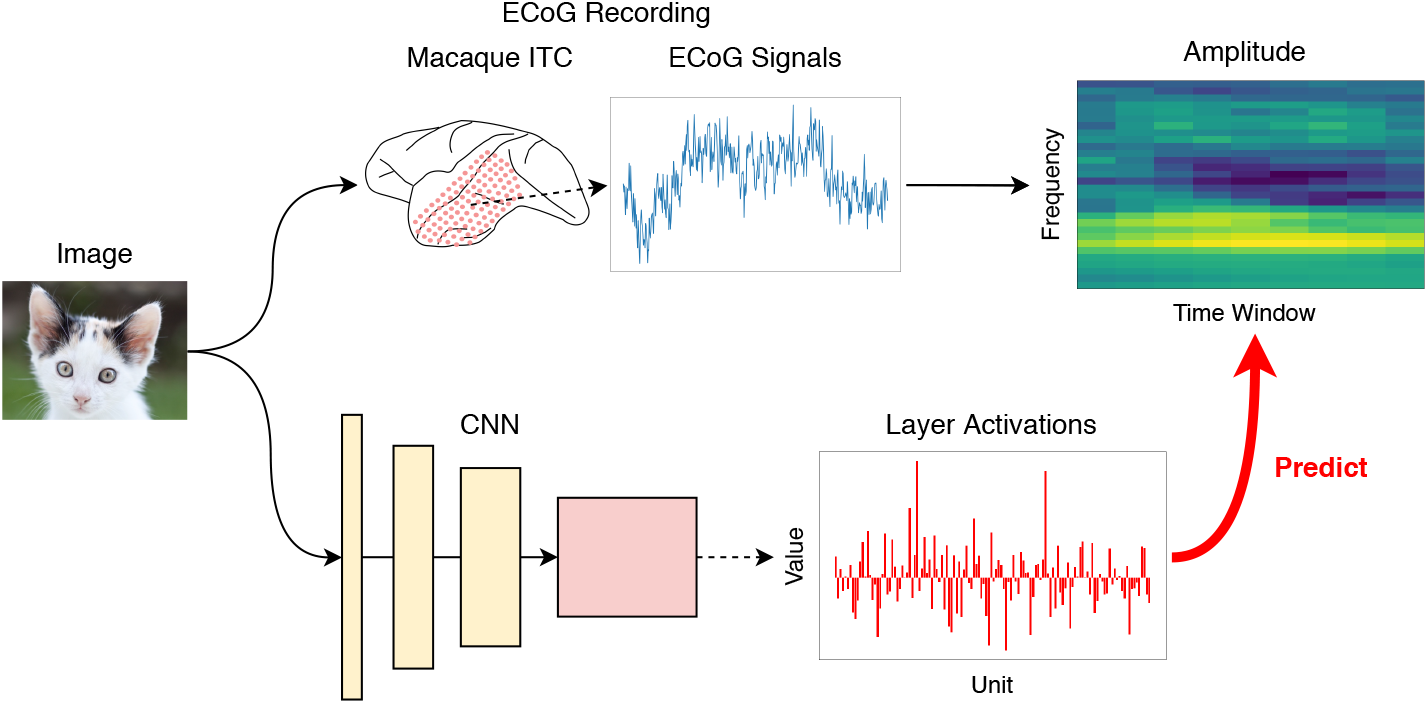
Encoding frequency-specific ECoG activities from CNN features. We trained encoding models that predict frequency-specific ECoG activities given visual features extracted from a pretrained CNN. We recorded ECoG signals from the macaque ITC while presenting natural images, and extracted frequency-specific amplitude using time-frequency decomposition. Using the same image set, we extracted visual features from each convolution and fully-connected (FC) layer of a pretrained CNN. For convolutional layers, which have three dimensions (width, height, channels), we downsampled features over the width and height (global average pooling). After feature extraction, we trained ridge regression models that predict ECoG amplitude at a specific site, frequency, and time window from CNN features at a specific layer.

## 2 Materials and methods

### 2.1 Image set

We prepared diverse natural images from six object classes: building, body part, face, fruit, insect, and tool. First, 60,000 (10,000 per class) candidate images were selected by keyword search on *Flickr*. These images were then screened by a web-based survey using *Amazon Mechanical Turk*. For each class, the participant was given a brief instruction on image selection (mostly regarding image quality), and was presented six good and three bad example images. Each candidate image was evaluated by three participants. Images selected by all the three participants were considered as verified. For each class, if more than 1,000 images were verified, randomly-chosen 1,000 images among verified ones were used for our experiment. For stimulus presentation, original images were cropped to 512 × 512 pixels.

### 2.2 ECoG recording

We recorded ECoG signals from two female macaques (Macaca fuscata, monkey 1: 6.1 kg, monkey 2: 5.1 kg). All animal procedures were complied with the National Institute of Health Guide for the Care and Use of Laboratory Animals, and the Guide of the National BioResource Project “Japanese Monkeys” of the Ministry of Education, Culture, Sports and Technology (MEXT), Japan. The Niigata University Institutional Animal Care and Use Committee approved the experimental protocols.

For recording ECoG signals from the macaque inferior temporal cortex (ITC), we designed a 128-channel electrode grid that covers an area of 20 mm × 40 mm with a inter-grid distance of 2.5 mm. The electrode was fabricated on a 20 μm-thick flexible Parylene-C film using micro-electro-mechanical systems technology. One side of the each square contact was 1 mm.

The ECoG electrode was subdurally implanted under an aseptic conditions. After premedication with ketamine (50 mg/kg) and medetomidine (0.03 mg/kg), each monkey was intubated with an endotracheal tube of 6 or 6.5 mm and connected to an artificial respirator (A.D.S. 1,000, Engler engineering corp., FL, USA). The venous line was secured using lactated Ringer’s solution, and ceftriaxone (100 mg/kg) was dripped as a prophylactic antibiotic. The body temperature was maintained to keep around 37 °C using an electric heating mat. A vacuum fixing bed (Vacuform, B.u.W.Schmidt GmbH, Garbsen, Germany) was used to maintain the position of the body. The oxygen saturation, heart rate, and end-tidal CO2 were continuously monitored (Surgi Vet, Smiths medical PM inc., London, UK) throughout the surgery to adjust the level of anesthesia. The skull was fixed with a 3-point fastening device (Integra Co., NJ, USA) with a custom-downsized attachment for macaques. The target location and the size of craniotomy were determined using preoperative magnetic resonance imaging. In the intra-dural operation, we used a microscope (Ophtalmo-Stativ S22, Carl Zeiss Inc., Oberkochen, Germany) with a CMOS color camera (TS-CA-130MIII, MeCan Imaging Inc., Saitama, Japan). The electrode grid was carefully attached onto the cortical surface, and the dura was closed with water tight suturing to prevent cerebrospinal fluid leakage. The electrode lead, microconnectors (Omnetics, MN, USA), and a custom-made plastic connector-chamber (Vivo, Hokkaido, Japan) were fixed onto the bone with resin.

The monkeys were trained with a visual fixation task to keep their gazes within ±1.5 degree of visual angle around the fixation target. Eye movements were captured with an infra-red camera system with a sampling rate of 60 Hz.

The stimuli were presented on a 15-inch CRT monitor (NEC, Tokyo, Japan) at a viewing distance of 26 cm. After 450 ms of stable fixation, the stimulus was presented for 300 ms followed by a 600-ms blank interval. In each trial, two-to-five stimuli were successively presented. The monkeys were rewarded with a drop of apple juice for maintaining their fixation over the entire duration of the trial. The long axis of each stimulus subtended six degree of visual angle.

Signals were differentially amplified using a 128-channel amplifier (Plexon, TX, USA or Tucker Davis Technologies, FL, USA) with high- and low-cutoff filters at 300 Hz and 1.0 Hz, respectively. All subdural electrodes were referenced to a titanium screw that was attached directly to the dura at the vertex area. Recording was conducted at a sampling rate of 1 kHz per channel.

### 2.3 Time-frequency decomposition

As preprocessing, we first eliminated line noise in raw ECoG signals by applying a third-order Butterworth filter at 50 Hz. Then, we rereferenced signals at all the channels by taking bipolar derivatives between neighboring channels. Because electric potentials recorded by ECoG often contain noise from a non-cortical reference site or non-local signals, this rereferencing procedure can help us extract more local electric activities on the cortical surface. In total, we used ECoG signals at 112 sites extracted from the original 128 channels.

We computed the analytic amplitude at 30 frequencies using complex Morlet wavelet convolution. The central frequencies were logarithmically sampled from 1 to 250 Hz. For each central frequency *f* (Hz), we constructed a complex Morlet wavelet:

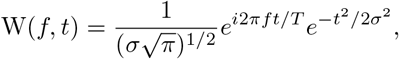

where *t* is a time step (ms), *σ* = *n*_*f*_/(2*πf*) is the standard deviation, and *n*_*f*_ is the number of wavelet cycles. The number of wavelet cycles *n*_*f*_ for each central frequency was logarithmically sampled from 3 to 14. We computed the analytic amplitude as the absolute value of the convolution results between preprocessed ECoG signals and the wavelet:

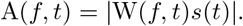

After extracting the analytic amplitude, we conducted postprocessing for coping with trial-by-trial and temporal differences. We first normalized each trial’s activities with the average amplitude in the baseline (−500 to −201 ms relative to the stimulus onset) using decibel conversion:

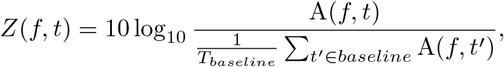

where *T*_*baseline*_ is the number of time steps in the baseline (300 ms). After baseline normalization, we took the cross-trial average of normalized decibel changes over five trials for each stimulus. Finally, we downsampled multi-trial activities in nine sliding time windows (1-100, 51-150, 101-200, 151-250, 201-300, 251-350, 301-400, 351-450, and 401-500 ms relative to the stimulus onset). As the result, we obtained frequency-specific ECoG activities for 112 sites, 30 central frequencies, and 9 time windows for each monkey.

### 2.4 Extracting hierarchical visual features from convolutional neural networks

Deep convolutional neural networks (CNNs) have achieved state-of-the-art performance on diverse computer vision tasks, such as object recognition, semantic segmentation, object detection, and video recognition [15, 16, 17, 18]. Several previous studies [22, 23, 24] showed that representations in pretrained CNNs are efficiently applicable on novel image sets and tasks (e.g., object category classification, scene recognition, fine grained recognition, attribute detection and image retrieval). Furthermore, CNNs enable us to extract hierarchical visual features, since CNNs have its layer hierarchy and previous studies indicate that lower, mid, and higher CNN layers represent lower-, mid-, and higher-level visual features, respectively [19]. Therefore, pretrained CNNs are useful for extracting optimized hierarchical visual features. Interestingly, several recent studies in cognitive neuroscience investigated brain activities using hierarchical visual features from CNNs, and found the similarity between the layer hierarchy of CNNs and the anatomical hierarchy of the primate ventral stream [20, 21].

In this work, we used a pretrained CNN for analysing frequency-specific ECoG activities. We employed a VGG-16 network [25] that was pretrained on the ILSVRC2012 object classification task [26]. We fed our image set to the pretrained CNN, and extracted visual features at each layer. VGG-16 consists of 13 convolution layers, 5 max pooling layers, 3 fully-connected (FC) layers, and the final classification (softmax) layer. We extracted visual features from all the convolution and FC layers, resulting in 16 layers used in total.

While features at FC layers are vectors, those at convolution layers are three-dimensional tensors that have width, height, and depth (channels). In our experiment, we downsampled the output of convolution layers by taking the spatial average (global average pooling) over the width and height.

### 2.5 Encoding ECoG features from CNN features

To compare the prediction performance over the frequency bands and CNN layers, we trained and evaluated encoding models that predict frequency-specific ECoG activities from visual features at a specific CNN layer. More specifically, each encoding model was trained to predict ECoG activities at a specific site, frequency, and time window, given CNN features at a specific layer as input:

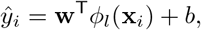

where **x**∈ ℝ^3×224×224^ is the image, 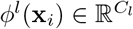 is a CNN feature at *l*-th layer, 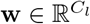 is a weight vector that projects CNN features into a scalar, and *b* ∈ ℝ is a bias term. The parameters, **w** and *b*, were optimized using ridge regression:

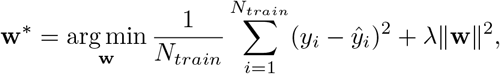

where *N*_*train*_ is the number of training samples, and λ is a hyperparameter to control the penalty term. Encoding models were independently trained for each combination of ECoG site, frequency, time window, and CNN layer.

We split the image set so that no image in the validation and test set is included in the training set. In monkey 1’s dataset, the training, validation, and test set contains 2431, 808, and 808 stimuli, respectively. In monkey 2’s dataset, the training, validation, and test set contains 2441, 813, and 813 stimuli, respec tively. Before training encoding models, CNN features in the training set were standardized so that they have zero mean and unit variance. CNN features in the validation and test sets were normalized with the mean and standard deviation in the training set. We implemented our encoding models with *PyTorch* [27]. We optimized our encoding models using the *Adam* optimizer [28], with a learning rate of 10^−4^, *β*_1_ = 0.9, *β*_2_ = 0.999, a weight decay (*λ*) of 10^−6^, and a batch size of 128. The maximum training epoch was 100 epochs, but we stopped training when the validation performance was not improved for successive 10 epochs.

In the test set, we evaluated each encoding model’s prediction performance as Pearson’s correlation coefficient between ground truth and predicted values. To eliminate results that can occur by chance, we determined the significance threshold of Pearson’s correlation coefficient using the permutation test. For each encoding model, we computed the correlation between ground truth and randomly-permuted predictions. We repeated the procedure for 1,000 permutations, and took the largest value as the threshold.

## 3 Results

### 3.1 Theta and gamma bands are better predicted from CNN features

First, we compare the difference of the prediction performance over the frequency bands. Comparing the prediction performance over the frequency bands shows us which frequency bands are more related to visual features extracted from a pretrained CNN. Our hypothesis here is that specific frequency bands show better performance than the other bands. That is, specific frequency bands might show stronger selectivity to visual features in CNNs.

Figure 2 shows the comparison of the prediction performance over the frequency bands. For each ECoG site, we took the maximum performance over the time windows and CNN layers for visualization. For both monkeys, the average prediction performance over the sites was peaked at theta (around 5 Hz) and gamma bands (around 20-25 Hz), showing that ECoG activities in the low- and high-frequency bands were better predicted from CNN features than the other frequency bands. In delta (1-2 Hz) and high-gamma bands (100-200 Hz), several sites showed the prediction performance around 0.3, but the curve of the mean prediction performance did not peak at these frequency bands.

**Figure 2:**
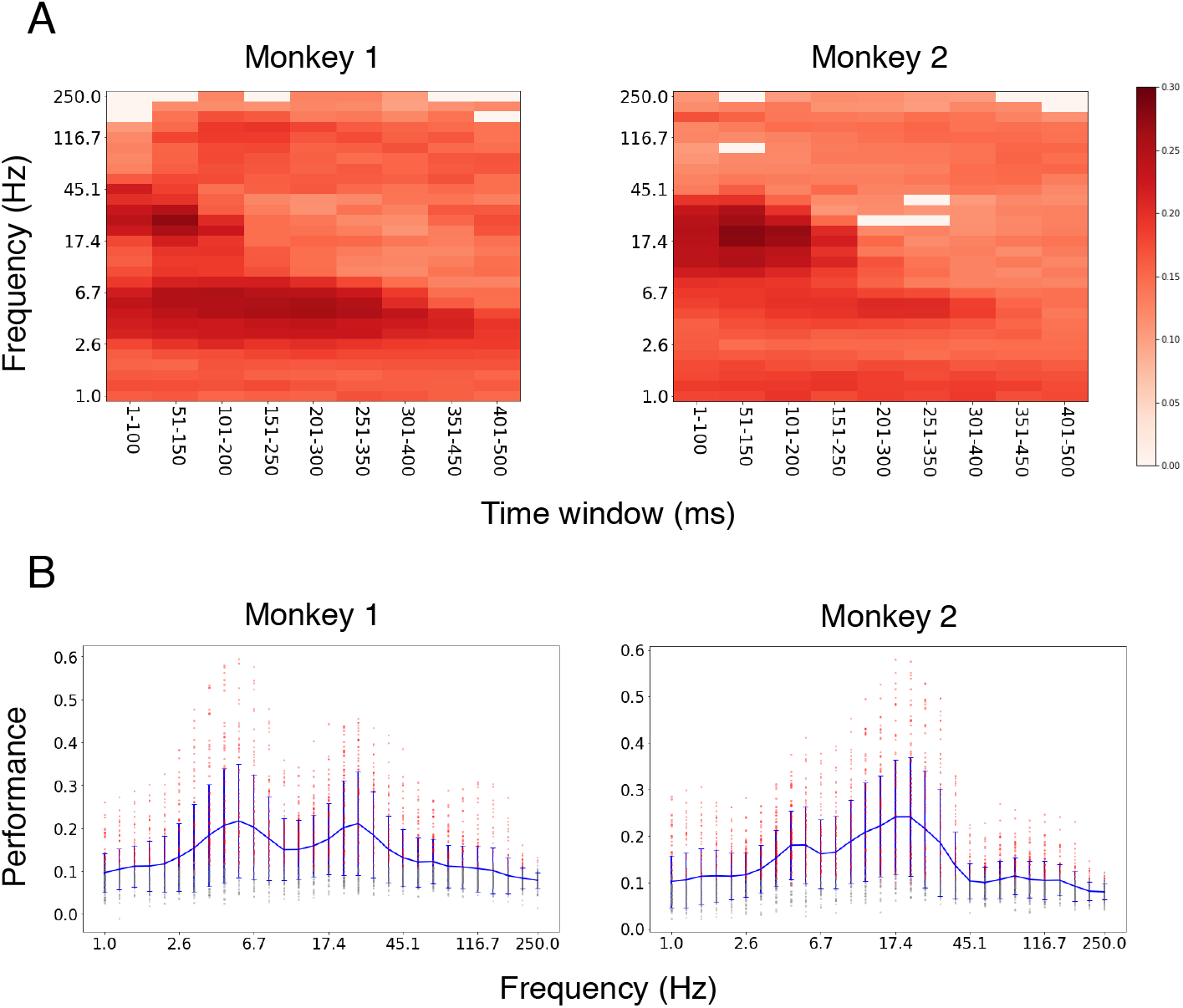
Comparison of the prediction performance. In the test set, the prediction performance was measured as Pearson’s correlation coefficient between ground truth and predicted values. (A) The prediction performance over the frequencies and time windows. For each site, the maximum performance over the CNN layers was extracted. The average performance over sites that showed better performance than the significance threshold (*p* < 0.0001 in the permutation test) is shown here. (B) The prediction performance over the frequencies. Red dots indicate the prediction performance of each ECoG site. For each site, the maximum performance over the time windows and CNN layers was extracted. Only results above the significance threshold are shown here (*p* < 0.0001 in the permutation test). Blue line indicate the mean prediction performance over the ECoG sites. Blue error bars indicate the standard error of the prediction performance over the ECoG sites.

### 3.2 Theta and gamma bands are better predicted from higher and lower CNN layers, respectively

In visual object recognition, CNNs receive input images, perform convolution and subsampling (pooling) operations at each layer, and finally output object-level classification results. Therefore, units in higher CNN layers have larger receptive fields. Furthermore, several previous studies analyzed the visual selectivity of units in each layer, and showed that lower and higher layers have selectivity to lower- (e.g., color, orientation, grating, texture) and higher-level (e.g., shape, object part, animal face) visual features [19, 29]. Since visual features in higher CNN layers have larger receptive fields and are correlated with higher-level visual patterns, we can investigate the difference of theta- and gamma-band activities in terms of the hierarchy of visual features in CNNs. Here, we visualized the relationship between the CNN layers and the frequency bands.

Figure 3 shows the comparison of layer assignments for theta and gamma bands. For both monkeys, in gamma band, most sites were assigned lower convolution layers. On the other hand, in theta band, most sites were assigned higher convolution and FC layers. Several sites in theta band were assigned the highest layer (FC8), which outputs class-wise logits for classification, but no site in gamma band was assigned the FC8 layer. These results show the distinct visual selectivity of theta- and gamma-band ECoG activities in terms of the layer hierarchy of CNNs.

**Figure 3:**
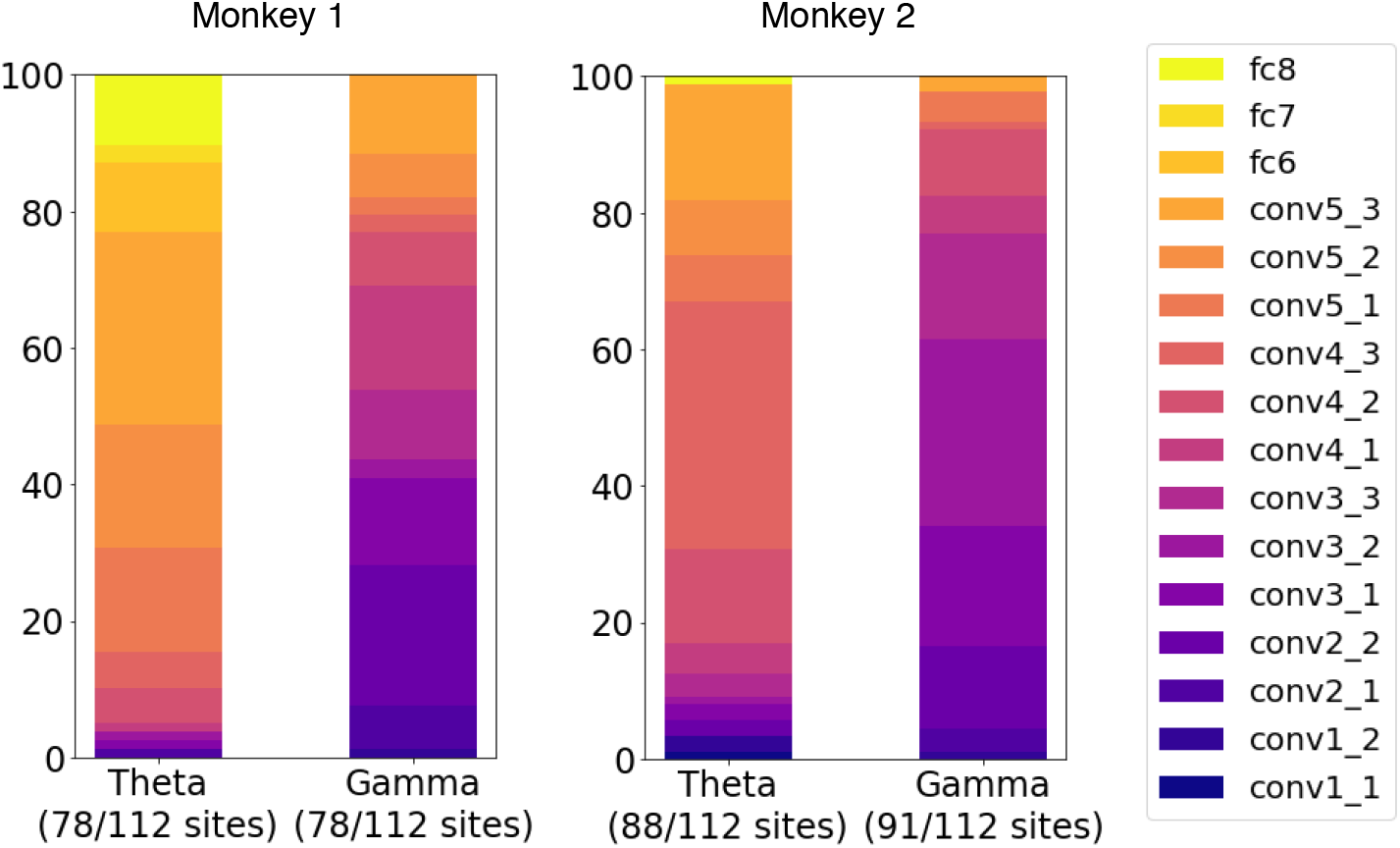
Comparing the proportion of assigned CNN layers for each frequency. For each site, the maximum performance over the time windows is extracted. Then, comparing the prediction performance over the CNN layers, the best layer is assigned to the site if the prediction performance is better than the significance threshold (*p* < 0.0001 in the permutation test). The vertical axis shows the proportion of each layer. The numbers in the parentheses show the number of sites above the threshold.

### 3.3 Visualizing the selectivity of theta- and gamma-band encoding models

Next, we examined more detailed visual selectivity of theta and gamma bands using optimized encoding models. Each encoding model first transforms an image with convolution and pooling operations in the pretrained CNN, and then projects CNN features into frequency-specific prediction value. Starting from random images, we can optimize the image so as to maximize the prediction value with gradient descent [30, 31]:

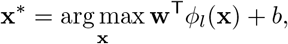

where *ϕ*_*l*_ is a feature extractor at a specific CNN layer given an image. On the other hand, in minimization, we optimize the image so as to minimize the prediction value:

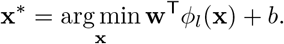

With this method, we can qualitatively investigate what kind of visual patterns are preferred by theta- and gamma-band encoding models. Using the pretrained CNN and frequency-specific encoding models, we optimize a randomly-initialized image using the Adam optimizer with a learning rate of 0.01, *β*_1_ = 0.9, and *β*_2_ = 0.999. To avoid noisy results, we stochastically transformed the image before feeding into the CNN [31]. More specifically, we stochastically jittered (0-8 pixels), rotated (0-45 degree), and scaled (1.0-1.8 times) the image. We also extracted nine top and bottom images based on test-set predictions of each encoding model.

Several examples of optimized and preferred images are shown in Figure 4. For theta-band models, preferred images appeared to be strongly related to higher-level visual features (e.g., face), and optimized images tended to contain higher-level, more complex visual patterns. In contrast, for gamma-band models, preferred images appeared to be related to lower-level visual features, and optimized images tended to contain lower-level, more local visual patterns.

**Figure 4:**
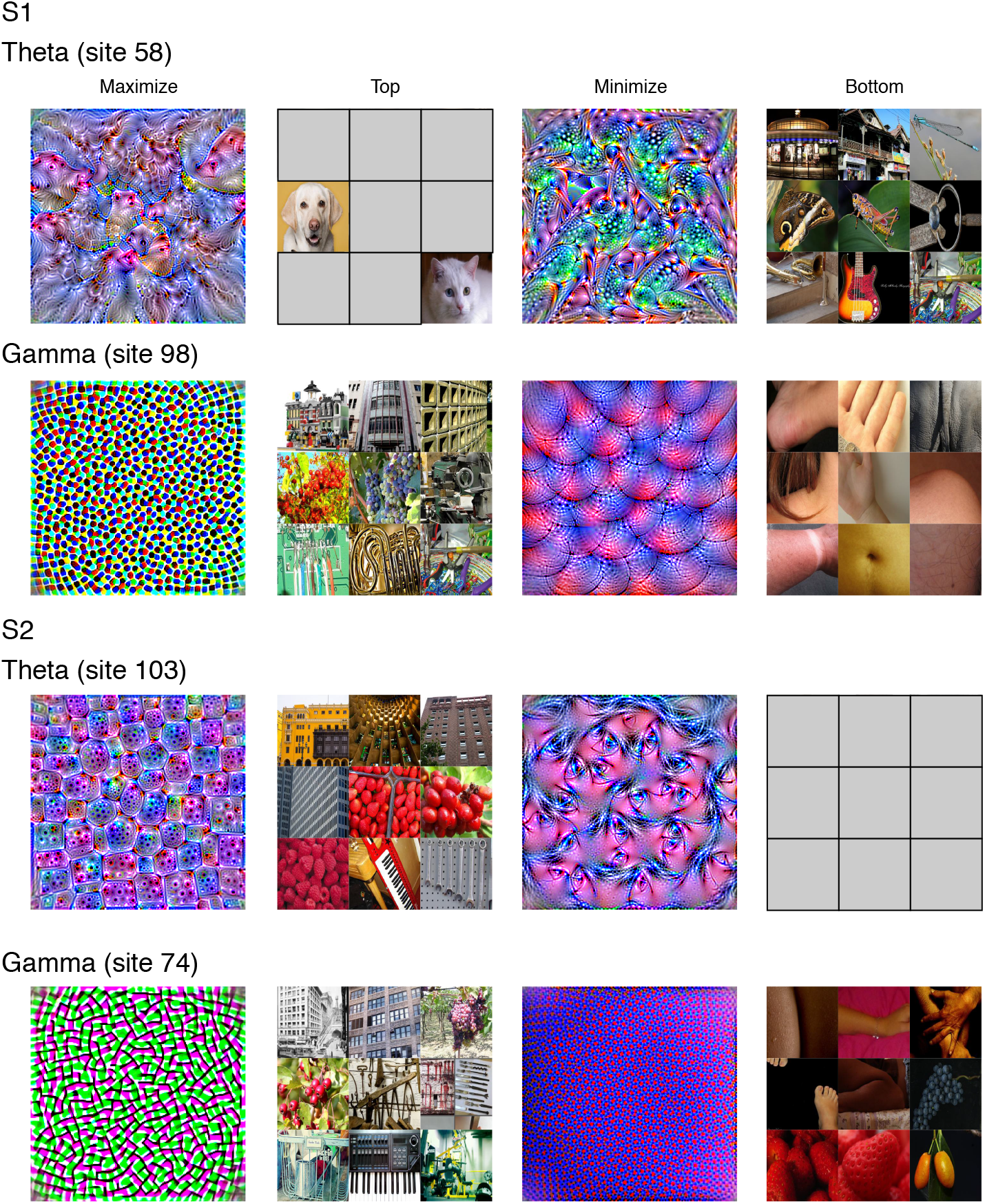
Examples of optimized (maximize, minimize) and preferred (top, bottom) images for theta- and gamma-band encoding models. Optimized images were produced by updating randomly-initialized images so as to maximize or minimize the predicted value of each encoding model. Preferred images were selected based on predicted values of each encoding model on the test set. Face images were shown as grey images to exclude any identifying information of people.

## 4 Discussion

In this work, we have analyzed frequency-specific ECoG activities in the macaque ITC using hierarchical visual features extracted from a pretrained CNN. By analyzing the prediction performance and visual selectivity of our encoding models, we were able to quantitatively and qualitatively investigate the frequency-specific visual selectivity of ECoG activities. Comparing the prediction performance of our encoding models, we observed that ECoG activities in two time-frequency bands, theta and gamma bands, were better predicted from CNN features than the other bands. In the comparison of assigned CNN layers, we found that theta- and gamma-band activities were better predicted from higher and lower layers, respectively. Furthermore, our visualization analysis qualitatively showed that theta- and gamma-band activities were related to higher- and lower-level visual features, respectively. Our results indicate that distinct levels of visual information are multiplexed in low- and high-frequency activities in brain signals.

There have been several previous studies on the visual selectivity of frequency-specific neuronal activities in the visual cortex. For example, Belitski *et al.* [8] analyzed local field potentials (LFPs) and multi-unit activities (MUAs) recorded from the primary visual cortex of anaesthetized monkeys. Their results indicate that: (1) lower (< 12 Hz) and high-gamma (60-100 Hz) bands are more related to visual information, and (2) these two bands are not correlated with each other, implying that these two time-frequency bands represent distinct visual features. Similar results were also observed in [9], where human EEG activities in multiple time-frequency bands are analyzed on a face image set. They found that lower- and higher-frequency bands were related to distinct facial features. Jacobs *et al.* [6] analyzed ECoG signals recorded from neurosurgical patients. Their results indicate that: (1) gamma band (25-128 Hz) is more informative than the other bands for classifying letter identities, and (2) gamma band has strong phase-amplitude coupling with theta band (4-8 Hz). Recently, similar to our work, Kuzovkin *et al.* [32] analyzed frequency-specific ECoG activities in the human visual cortex using a pretrained CNN. They employed the *representation similarity analysis* (RSA) [33] to compare CNN layers and frequency-specific ECoG activities. They observed that, the layer hierarchy of CNNs matched the anatomical hierarchy of the human ventral visual pathway for gamma-band (30-150 Hz) ECoG activities. In this work, using hierarchical representations extracted from CNNs, we observed that theta- and gamma-band activities were more related to visual features than the other bands, and that these two time-frequency bands have distinct visual selectivity. It is noteworthy that we found the distinct visual selectivity of theta and gamma band activities on a more diverse set of natural images.

Our findings are related to recent studies on frequency-specific roles in feed-forward and feedback processing in the primate visual cortex. Bastos *et al.* [12] analyzed ECoG signals recorded from rhesus monkeys to investigate whether inter-areal influences between visual cortical areas are subserved differentially by different frequency bands. Their results indicate that feedforward influences are related to theta (~4 Hz) and gamma band (~60-80 Hz), and that feedback influences by beta band (~14-18 Hz). Similarly, van Kerkoerle *et al.* [34] reported that, in the monkey visual cortex, alpha (5-15 Hz) and gamma (40-90 Hz) bands are related feedback and feedforward processing between V1 and V4. These studies indicate that several different frequency bands are related to frequency-specific roles in feedforward or feedback processing between visual cortical areas. Combining our results with these previous studies on frequency-specific roles in inter-areal communications, it could be possible that theta and gamma bands carry distinct visual information for different roles in inter-areal communications in the visual cortex.

## Acknowledgements

This work was supported by JSPS KAKENHI Grant number JP15H05919 and 16K01959. H. D. was supported by the doctoral course scholarship of Division for Interdisciplinary Advanced Research and Education (DIARE), Tohoku University.

## Author contributions

D., K. K., and T. O. designed the study. K. K. and I. H. conducted ECoG recording. H. D. developed, implemented, and conducted computational experiments. H. D., K. K., and T. O. wrote the paper.

## References

[1] Andrew J Watrous, Juergen Fell, Arne D Ekstrom, and Nikolai Axmacher. More than spikes: common oscillatory mechanisms for content specific neural representations during perception and memory. Current opinion in neurobiology, 31:33–39, 2015.

[2] Pascal Fries. Rhythms for cognition: communication through coherence. Neuron, 88(1):220–235, 2015.

[3] György Buzsáki and Andreas Draguhn. Neuronal oscillations in cortical networks. Science, 304(5679):1926–1929, 2004.

[4] Christoph Kayser and Peter König. Stimulus locking and feature selectivity prevail in complementary frequency ranges of v1 local field potentials. European Journal of Neuroscience, 19(2):485–489, 2004.

[5] Philipp Berens, Georgios A Keliris, Alexander S Ecker, Nikos K Logothetis, and Andreas S Tolias. Feature selectivity of the gamma-band of the local field potential in primate primary visual cortex. Frontiers in Neuroscience, 2:37, 2008.

[6] Joshua Jacobs and Michael J Kahana. Neural representations of individual stimuli in humans revealed by gamma-band electrocorticographic activity. Journal of neuroscience, 29(33):10203–10214, 2009.

[7] Christopher M Lewis, Conrado A Bosman, Nicolas M Brunet, Bruss Lima, Mark J Roberts, Thilo Womelsdorf, Peter de Weerd, Sergio Neuenschwander, Wolf Singer, and Pascal Fries. Two frequency bands contain the most stimulus-related information in visual cortex. bioRxiv, page 049718, 2016.

[8] A. Belitski, A. Gretton, C. Magri, Y. Murayama, M. A. Montemurro, N. K. Logothetis, and S. Panzeri. Low-frequency local field potentials and spikes in primary visual cortex convey independent visual information. Journal of Neuroscience, 28(22):5696–5709, 2008.

[9] Philippe G Schyns, Gregor Thut, and Joachim Gross. Cracking the code of oscillatory activity. PLoS Biology, 9(5):e, 2011.

[10] Siddhesh Salelkar, Gowri Manohari Somasekhar, and Supratim Ray. Distinct frequency bands in the local field potential are differently tuned to stimulus drift rate. Journal of Neurophysiology, 120(2):681–692, 2018.

[11] Astrid Von Stein, Carl Chiang, and Peter König. Top-down processing mediated by interareal synchronization. Proceedings of the National Academy of Sciences, 97(26):14748–14753, 2000.

[12] Andre Moraes Bastos, Julien Vezoli, Conrado Arturo Bosman, Jan-Mathijs Schoffelen, Robert Oostenveld, Jarrod Robert Dowdall, Peter De Weerd, Henry Kennedy, and Pascal Fries. Visual areas exert feedforward and feedback influences through distinct frequency channels. Neuron, 85(2):390–401, 2015.

[13] Georgios Michalareas, Julien Vezoli, Stan Van Pelt, Jan-Mathijs Schoffelen, Henry Kennedy, and Pascal Fries. Alpha-beta and gamma rhythms subserve feedback and feedforward influences among human visual cortical areas. Neuron, 89(2):384–397, 2016.

[14] Craig G Richter, William H Thompson, Conrado A Bosman, and Pascal Fries. Top-down beta enhances bottom-up gamma. Journal of Neuroscience, 37(28):6698–6711, 2017.

[15] Alex Krizhevsky, Ilya Sutskever, and Geoffrey E Hinton. Imagenet classification with deep convolutional neural networks. In Advances in Neural Information Processing Systems, pages 1097–1105, 2012.

[16] Kaiming He, Xiangyu Zhang, Shaoqing Ren, and Jian Sun. Delving deep into rectifiers: Surpassing human-level performance on imagenet classification. In Proceedings of the IEEE International Conference on Computer Vision, pages 1026–1034, 2015.

[17] Jonathan Long, Evan Shelhamer, and Trevor Darrell. Fully convolutional networks for semantic segmentation. In Proceedings of the IEEE Conference on Computer Vision and Pattern Recognition, pages 3431–3440, 2015.

[18] Oriol Vinyals, Alexander Toshev, Samy Bengio, and Dumitru Erhan. Show and tell: A neural image caption generator. In Proceedings of the IEEE Conference on Computer Vision and Pattern Recognition, pages 3156–3164, 2015.

[19] Matthew D Zeiler and Rob Fergus. Visualizing and understanding convolutional networks. In European Conference on Computer Vision, pages 818–833. Springer, 2014.

[20] Daniel LK Yamins, Ha Hong, Charles F Cadieu, Ethan A Solomon, Darren Seibert, and James J DiCarlo. Performance-optimized hierarchical models predict neural responses in higher visual cortex. Proceedings of the National Academy of Sciences, 111(23):8619–8624, 2014.

[21] Umut Güçlü and Marcel AJ van Gerven. Deep neural networks reveal a gradient in the complexity of neural representations across the ventral stream. Journal of Neuroscience, 35(27):10005–10014, 2015.

[22] Jason Yosinski, Jeff Clune, Yoshua Bengio, and Hod Lipson. How transferable are features in deep neural networks? I. Advances in Neural Information Processing Systems, pages 3320–3328, 2014.

[23] Ali Sharif Razavian, Hossein Azizpour, Josephine Sullivan, and Stefan Carlsson. Cnn features off-the-shelf: an astounding baseline for recognition. In Proceedings of the IEEE Conference on Computer Vision and Pattern Recognition Workshops, pages 806–813, 2014.

[24] Jeff Donahue, Yangqing Jia, Oriol Vinyals, Judy Hoffman, Ning Zhang, Eric Tzeng, and Trevor Darrell. Decaf: A deep convolutional activation feature for generic visual recognition. In ICML, pages 647–655, 2014.

[25] Karen Simonyan and Andrew Zisserman. Very deep convolutional networks for large-scale image recognition. arXiv preprint arXiv:1409.1556, 2014.

[26] Olga Russakovsky, Jia Deng, Hao Su, Jonathan Krause, Sanjeev Satheesh, Sean Ma, Zhiheng Huang, Andrej Karpathy, Aditya Khosla, Michael Bernstein, Alexander C. Berg, and Li Fei-Fei. ImageNet Large Scale Visual Recognition Challenge. International Journal of Computer Vision (IJCV), 115(3):211–252, 2015.

[27] Adam Paszke, Sam Gross, Francisco Massa, Adam Lerer, James Bradbury, Gregory Chanan, Trevor Killeen, Zeming Lin, Natalia Gimelshein, Luca Antiga, et al. Pytorch: An imperative style, high-performance deep learning library. In Advances in Neural Information Processing Systems, pages 8024–8035, 2019.

[28] Diederik P Kingma and Jimmy Ba. Adam: A method for stochastic optimization. arXiv preprint arXiv:1412.6980, 2014.

[29] David Bau, Bolei Zhou, Aditya Khosla, Aude Oliva, and Antonio Torralba. Network dissection: Quantifying interpretability of deep visual representations. In Proceedings of the IEEE Conference on Computer Vision and Pattern Recognition, pages 6541–6549, 2017.

[30] Dumitru Erhan, Yoshua Bengio, Aaron Courville, and Pascal Vincent. Visualizing higher-layer features of a deep network. University of Montreal, 1341(3):1, 2009.

[31] Chris Olah, Alexander Mordvintsev, and Ludwig Schubert. Feature visualization. Distill, 2(11):e, 2017.

[32] Ilya Kuzovkin, Raul Vicente, Mathilde Petton, Jean-Philippe Lachaux, Monica Baciu, Philippe Kahane, Sylvain Rheims, Juan R Vidal, and Jaan Aru. Activations of deep convolutional neural networks are aligned with gamma band activity of human visual cortex. Communications biology, 1(1):107, 2018.

[33] Nikolaus Kriegeskorte, Marieke Mur, and Peter A Bandettini. Representational similarity analysis-connecting the branches of systems neuroscience. Frontiers in systems neuroscience, 2:4, 2008.

[34] Timo Van Kerkoerle, Matthew W Self, Bruno Dagnino, Marie-Alice Gariel-Mathis, Jasper Poort, Chris Van Der Togt, and Pieter R Roelfsema. Alpha and gamma oscillations characterize feedback and feedforward processing in monkey visual cortex. Proceedings of the National Academy of Sciences, 111(40):14332–14341, 2014.

